# *Cis*-directed cleavage and nonstoichiometric abundances of 21-nt reproductive phasiRNAs in grasses

**DOI:** 10.1101/243907

**Authors:** Saleh Tamim, Zhaoxia Cai, Sandra Mathioni, Jixian Zhai, Chong Teng, Qifa Zhang, Blake C. Meyers

## Abstract

- Post-transcriptional gene silencing in plants results from independent activities of diverse small RNA types. In anthers of grasses, hundreds of loci yield non-coding RNAs that are processed into 21- and 24-nt phased small interfering RNAs (phasiRNAs); these are triggered by miR2118 and miR2275.
- We characterized these “reproductive phasiRNAs” from rice panicles and anthers across seven developmental stages. Our computational analysis identified characteristics of the 21-nt reproductive phasiRNAs that impact their biogenesis, stability, and potential functions.
- We demonstrate that 21-nt reproductive phasiRNAs can function in *cis* to target their own precursors. We observed evidence of this *cis* regulatory activity in both rice (*Oryza sativa*) and maize (*Zea mays*). We validated this activity with evidence of cleavage and a resulting shift in the pattern of phasiRNA production.
- We characterize biases in phasiRNA biogenesis, demonstrating that the Pol II-derived “top” strand phasiRNAs are consistently higher abundance than the bottom strand. The first phasiRNA from each precursor overlaps the miR2118 target site, and this impacts phasiRNA accumulation or stability, evident in the weak accumulation of this phasiRNA position. Additional influences on this first phasiRNA duplex include the sequence composition and length, and we show that these factors impact Argonaute loading.

## Introduction

Plants possess a wide variety of small RNAs (sRNAs) that play important roles during development (Chen 2009). sRNA classes specifically involved in post-transcriptional RNA silencing are microRNAs (miRNAs) and secondary small interfering RNAs (siRNAs) (Jones-Rhoades *et al*. 2006), distinguished mainly by their biogenesis. Secondary siRNAs require a miRNA trigger to direct cleavage of their RNA precursor that is subsequently converted to double-stranded RNA (dsRNA) by RNA DEPENDENT RNA POLYMERASE 6 (RDR6). miRNAs, transcribed initially as single-stranded mRNA precursors, fold to form a hairpin loop-like structure that is further processed to yield a small RNA duplex (Chen 2009, Jones-Rhoades *et al*. 2006).

Via advancements in DNA sequencing technology, the enormous growth in sRNA sequenced data sets coupled with genomic and mechanistic analyses has helped to further categorize siRNAs based on their size, origin, and/or known pathway information. *Trans*-acting siRNAs (tasiRNAs), a special class of phased siRNAs (phasiRNAs), are produced from their respective precursors, *TAS* loci, in a 21-nt “phasing” pattern because of processing by DICER-LIKE4 (DCL4). Four *TAS* loci (*TAS1/2/3/4*), first identified in Arabidopsis, all originate from non-coding transcripts. With the exception of the TAS3 locus, which has two target sites for the 21-nt miR390, Arabidopsis *TAS* loci are triggered by a 22-nt miRNA targeting a single site: *TAS1* and *TAS2* are triggered by miR173 and *TAS4* is triggered by miR828 (Allen *et al*. 2005, Axtell *et al*. 2006, Montgomery *et al*. 2008, Peragine *et al*. 2004, Rajagopalan *et al*. 2006, Vazquez *et al*. 2004, Yoshikawa *et al*. 2005). PhasiRNAs are the broader class of tasiRNA-like siRNAs, triggered by diverse miRNAs, originating from both coding and non-coding transcripts; some phasiRNAs function in *trans*, some in *cis*, and others may have no functionally relevant targets (Zhai *et al*. 2011). PhasiRNAs that are produced from protein-coding transcripts in many plant genomes are associated with genes modulating immunity and disease resistance (Fei et al., 2013). Diverse loci producing such disease-resistant phasiRNAs have been reported in a variety of eudicots, including Arabidopsis (Howell *et al*. 2007), apple (Xia *et al*. 2012), Medicago (Zhai *et al*. 2011), peach (Zhu *et al*. 2012), soybean (Arikit *et al*. 2014), and tomato (Shivaprasad *et al*. 2012).

In grass inflorescences, a high abundance of 21- and 24-nt phasiRNAs originate from non-coding transcripts, described in rice (Johnson *et al*. 2009, Song *et al*. 2012a), maize (Dukowic-Schulze *et al*. 2016, Zhai *et al*. 2015), and Brachypodium (Jeong *et al*. 2013). miR2118 and miR2275 trigger 21- and 24-nt reproductive phasiRNAs, respectively, across these species, and work in rice and maize demonstrates that these phasiRNAs and their triggers are particularly enriched in anthers, the male reproductive organ (Song *et al*. 2012a, Zhai *et al*. 2015). The pathway for the production of these reproductive phasiRNAs (hereafter called simply “phasiRNAs” for simplicity) has been studied in detail: DCL1 processes the miRNA triggers that directs cleavage of the *PHAS* precursor transcript; RDR6 and accessory proteins such as SUPPRESSOR OF GENE SILENCING 3 (SGS3) converts the precursor into dsRNA; a Dicer-like protein processes the dsRNA into phasiRNAs (DCL4 for 21-mers, DCL5 for 24-mers), starting from the miRNA target site; the resulting phasiRNAs are loaded into an ARGONAUTE (AGO) family protein to direct activity, typically silencing, of their corresponding targets (Borges and Martienssen 2015, Zhai *et al*. 2015). Work is still on-going to determine AGO specificity and functions, but in rice MEIOSIS ARRESTED AT LEPTOTENE 1 (MEL1), a homolog of Arabidopsis AGO5, loads 21-nt phasiRNAs (Komiya *et al*. 2014). Even though these phasiRNAs are numerous and highly abundant, their functions or activities have still not been investigated in much detail. Their localization and accumulation patterns are consistent with a role in reproductive biology, arguably analogous to animal PIWI-interacting RNAs (piRNAs), also produced from non-coding transcripts and are highly abundant in male organs (e.g. testis, for piRNAs)(Zhai *et al*. 2015). Recent work investigated methylation levels of maize *PHAS* loci and inferred that phasiRNAs could function in chromatin remodeling during chromosome pairing (Dukowic-Schulze *et al*. 2016). While the molecular basis of reproductive phasiRNA functions remain poorly characterized, at least the 21-nt class is clearly a component of full male fertility, as two such loci underlie photoperiod-sensitive male sterility in rice (Ding *et al*. 2012, Fan *et al*. 2016, Zhu and Deng 2012).

Plant reproductive phasiRNAs represent a rich source of data for studies of factors that influence their biogenesis, stability, and potential functions. In the species in which these phasiRNAs have been described, there are hundreds to thousands of coordinately expressed and processed loci, representing a large-scale dataset for analysis of these long, non-coding RNAs. In the current study, we sought to take advantage of this relatively untapped set of plant non-coding RNAs and phasiRNAs that they yield. To this end, we isolated and sequenced sRNAs from rice panicles across different stages of anther development, focusing on the premeiotic 21-nt phasiRNAs. We applied diverse computational analyses to these data, making comparisons with published maize sRNA data. Our analyses identified conserved patterns of 21-nt phasiRNA abundances, relative to the miR2118 trigger site. We show that individual phasiRNAs within a locus accumulate differentially; i.e. typically, one phasiRNA is highly abundant while others are absent. This observation holds also for MEL1-loaded 21-nt phasiRNAs. We also observed that the first-cycle phasiRNA is poorly represented among MEL1-associated small RNAs, and hence most likely not functional. Our analyses identified a number of factors that influence the representation of different phasiRNA positions, relative to the miRNA trigger site. Finally, we demonstrate that 21-nt reproductive phasiRNAs have *cis* regulatory activity, confirmed using Parallel Analysis of RNA Ends (PARE) data (German *et al*. 2009), and we model how these phasiRNAs function in *cis* to regulate their own transcripts.

## Materials and Methods

### Plant materials

The panicles and anthers used for library construction were collected from Nongken 58 (hereafter, “58N”), a *japonica* rice variety. Plants were grown under long-day conditions in the experimental farm of Huazhong Agricultural University (114.36°E, 30.48°N), Wuhan, China, as previous described (Ding *et al*. 2012).

### RNA extraction, library preparation and sequencing

Total RNA was isolated using TRIzol reagent (Invitrogen). Small RNA libraries were constructed using TruSeq Small RNA Sample Prep kits (Illumina, San Diego, CA). PARE libraries were constructed as previously described (Zhai *et al*. 2014). All libraries were sequenced on an Illumina HiSeq 2500 instrument at the Delaware Biotechnology Institute (Newark, DE).

### Bioinformatics analysis

sRNA and PARE raw data were processed by trimming the adapter sequence and removing low quality reads leaving reads that are 18-to 34-nt long (20-nt long for PARE reads). Processed reads were then mapped to the rice genome (*O. sativa* ssp. Japonica, MSU7) using Bowtie (Langmead *et al*. 2009) with no mismatches allowed. Twenty-one sRNA libraries and four PARE libraries from different anther developmental stages generated more than 450 million sRNA reads and 150 million PARE reads respectively (Supporting Information Table S1). For most of the comparisons across libraries, reads were normalized to transcripts per 10 million (TP10M), and reads mapping to more than one location (“hit”) in the genome were “hits-normalized” (the abundance is divided by the number of locations to which the read maps). sRNA reads for each developmental stage were merged together and the identification of *PHAS* loci (Supporting Information Table S2) was performed as previously described (Xia *et al*. 2013), requiring at least 10 unique reads to map into the identified locus (n ≥ 10) and at least 3 unique reads to map into a register (k ≥ 3). miR2118 targets of these loci were identified using TargetFinder (Fahlgren and Carrington 2010) using an optimal search algorithm (Smith and Waterman 1981) and with positions 2-13 receiving the same penalty score as the rest of the target positions. Sequence logos were generated using WebLogo 3.5 (Crooks *et al*. 2004) and most of the graphs were generated in R using ggplot package (Ginestet 2011). Statistical tests (t-test) were conducted using R. Images of individual *PHAS* loci were captured from the customized browser of the Meyers lab website (http://mpss.danforthcenter.org).

### Sequence Data

The small RNA and PARE data have been deposited to Gene Expression Omnibus (GEO) database with accession number GSE108238. We also used previously published maize and Arabidopsis data from GEO (GSE52293, GSE52297, and GSE35562) and rice MEL1 data from DNA Data Bank of Japan (DDBJ) with accession code DRP000161.

## Results

### Temporal regulation of reproductive phasiRNAs in rice

To examine 21-nt reproductive phasiRNAs in rice, we performed sRNA sequencing of developmentally staged young panicles and anthers in rice variety 58N. (Fig. 1a). We collected four stages of young panicles and three stages of anthers and generated three biological replicates for each stage for sequencing. After processing the sequenced reads (Supporting Information Table S1) and performing p-value based phasing analysis as previously described (Xia *et al*. 2013), 2107 21-nt *PHAS* loci and 57 24-nt *PHAS* loci were detected (Fig. 1b, Table S2). Both 21- and 24-nt phasiRNAs as well as their triggers (miR2118 and miR2275 respectively) exhibited temporal regulation in 58N (Fig. 1c), similar to what has been previously reported in maize (Zhai *et al*. 2015). Few phasiRNAs and miRNA triggers were detected in 0.1 cm panicles when the secondary branch primordium differentiates.

**Fig. 1.**
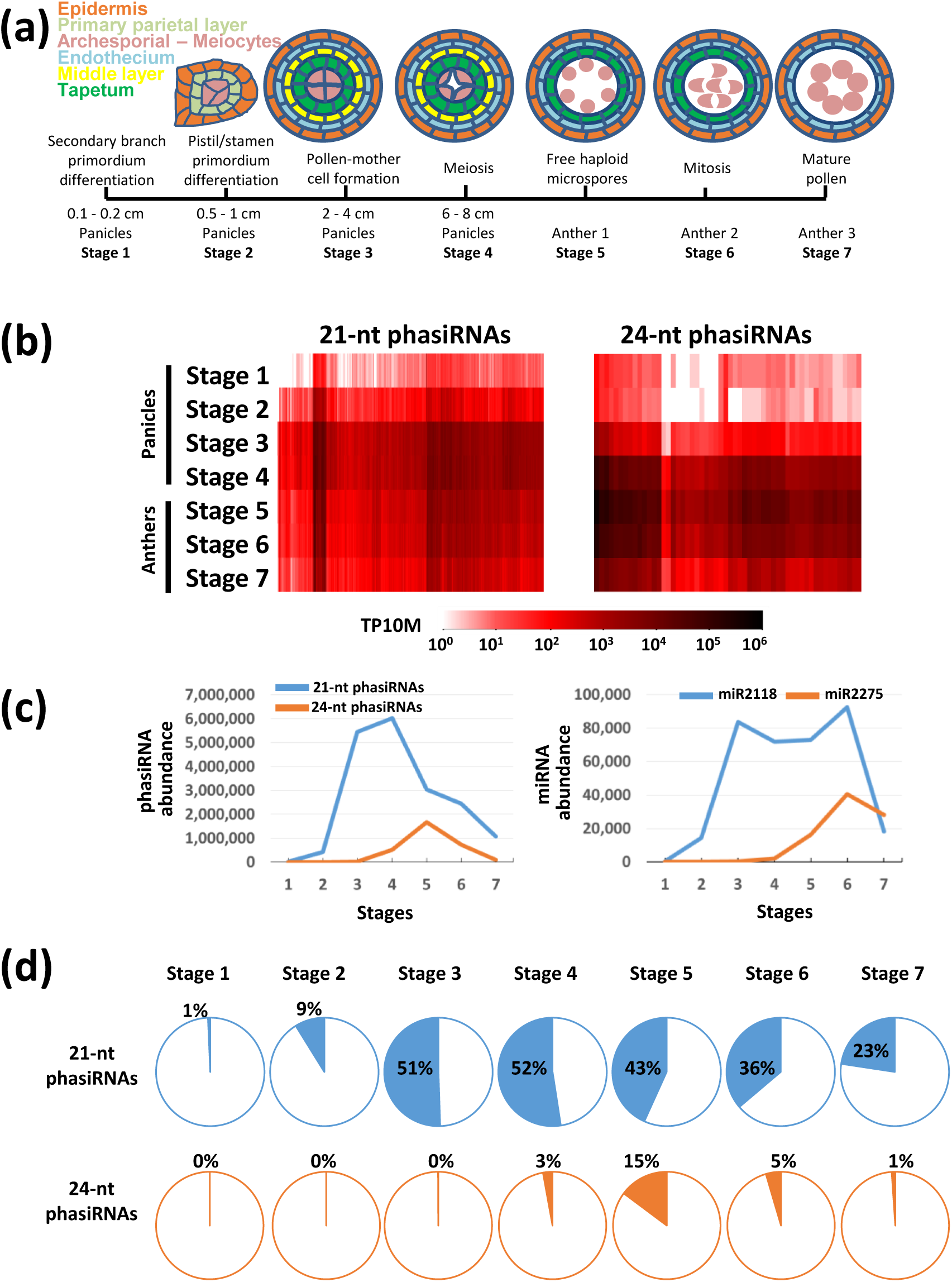
Reproductive phasiRNAs in rice exhibit temporal patterns of accumulation. (a) A schematic of cell patterns in a single lobe of a four-lobed grass anther, demonstrating the staging of the seven different points at which rice anthers were harvested for sequencing. Cell types indicated in colors shown in the key (upper left). (b) Heat maps depicting the abundances of 21-nt or 24-nt phased small interfering RNAs (phasiRNAs) at each stage (rows, as indicated) in rice (Nongken 58 variety under long day conditions); vertical bars indicate at left, 2107 21-nt loci, or at right, 57 24-nt loci. Abundances in both panels b and c are represented as transcripts per 10 million (TP10M), hit-normalized (see Methods). (c) Summed abundances of 21-nt or 24-nt phasiRNAs (left; summed data for all loci from panel b) and their microRNA (miRNA) triggers, miR2118 and miR2275, across the seven developmental stages. (d) Proportion of total 21-nt small RNAs (sRNAs) (top row) or total 24-nt sRNAs (bottom row) that are either 21-nt or 24-nt phasiRNAs, at each stage in rice. 21-nt phasiRNAs and their trigger miR2118 are represented with blue color while 24-nt phasiRNAs and their trigger miR2275 are represented in orange color.

The 21-nt phasiRNAs increase their accumulation at Stage 2 panicles (Fig. 1b), when the pistil/stamen primordia differentiate. The 21-nt phasiRNAs peaked at Stage 3 and 4 panicles (Fig. 1b), coincident with the progression from pollen-mother cell to meiosis, and when somatic cells continue differentiating out of their premeiotic supporting roles; at this peak, 21-nt phasiRNAs reached 52% of all 21-nt RNAs (Fig. 1d). This compares to 74% of all 21-mers at the peak in rice spikelets (Fei *et al*. 2016) and 60% in maize anther (Zhai *et al*. 2015); this lower proportion in rice panicles versus rice spikelets is likely just a dilution effect, as the panicle libraries included primary and secondary branches not known to generate phasiRNAs. Most 21-nt phasiRNAs were present until the mature pollen stage (Fig. 1b), but the abundance and proportion declined steadily after Stage 4 (Fig. 1c), at which point a large proportion of anthers have moved into meiosis. In contrast, most 24-nt phasiRNAs are undetectable until Stage 3 (Fig. 1b), at which point the panicles contain a high proportion of anthers forming pollen mother cells. The 24-nt phasiRNAs peaked at Stage 5, when meiosis finishes and free haploid microspores are released from the tetrads; at this peak, 24-nt phasiRNAs reached 15% of all 24-nt RNAs (Fig. 1d). This compares to 64% of the total 24-mers in isolated maize anthers (Zhai *et al*. 2015); this much lower proportion in rice may reflect the fact that there are fewer loci in rice compared to maize (57 vs 176, for the genotypes and stages analyzed). Alternatively, our sampling may have missed an even higher, narrow peak either side of Stage 5.

The abundance of the miR2118 family rose at Stage 2 panicles, peaked first at Stage 3, and stayed high through a second peak at Stage 6 anthers, and declined in mature pollen (Fig. 1c). Concurrently, the abundance of the miR2275 family members rose from Stage 4 to a peak in Stage 6, dropping off slightly in mature pollen (Fig. 1c). Unlike maize (Zhai *et al*. 2015), and perhaps due to the use of panicles instead of isolated anthers, we observed that both miR2118 and miR2275 abundances peaked after the abundance of the phasiRNA classes that they trigger.

### 21-nt reproductive phasiRNAs can function in *cis* to direct cleavage of their own precursors

To analyze 21-nt reproductive phasiRNAs in rice, we focused on the stages in which they are highly abundant, 6 to 8 cm panicles (Fig. 1c). We used the 2107 *PHAS* rice loci that we identified, together with previously-identified *PHAS* loci in maize (Zhai *et al*. 2015) for all subsequent analyses. We measured phasiRNA abundances (hits-normalized, see Methods) of each *PHAS* locus within the twenty registers (Axtell 2010, Axtell *et al*. 2006); for each register, we extended the analysis in the 3’ direction for twenty total phasiRNA duplexes (or “cycles”). The most abundant phasiRNAs appeared in register 0 (Fig. 2a,b; cycles and registers are explained in detail in Supporting Information Fig. S1), as expected and as previously reported (Johnson *et al*. 2009) since register 0 corresponds to phasiRNA processing initiated by miR2118-directed cleavage. However, phasiRNAs were also abundant in both species in registers 1, 9, 12, and 20 (Fig. 2a,b). Register 1 could be explained to result from either occasional DCL2 processing, since previous work demonstrated an accumulation of 22-nt siRNAs in *dcl4* (Xie *et al*. 2005), or slippage by DCL4 to process 22-nt phasiRNAs, which is known for Arabidopsis tasiRNAs (Chen *et al*. 2010). Either case would create a single register shift from the miR2118 target site. Additional, accumulating shifts, each at a low frequency, over the length of the phasiRNA would yield the observed decreasing abundances from register 1 to register 8. Register 20 similarly could result from DCL4 processing in inconsistent lengths, in this case shorter, as a single, 5’ register shift, a behavior known for Dicer proteins (Zhang *et al*. 2002). However, the peaks at register 9 and 12 were unexpected.

**Fig. 2.**
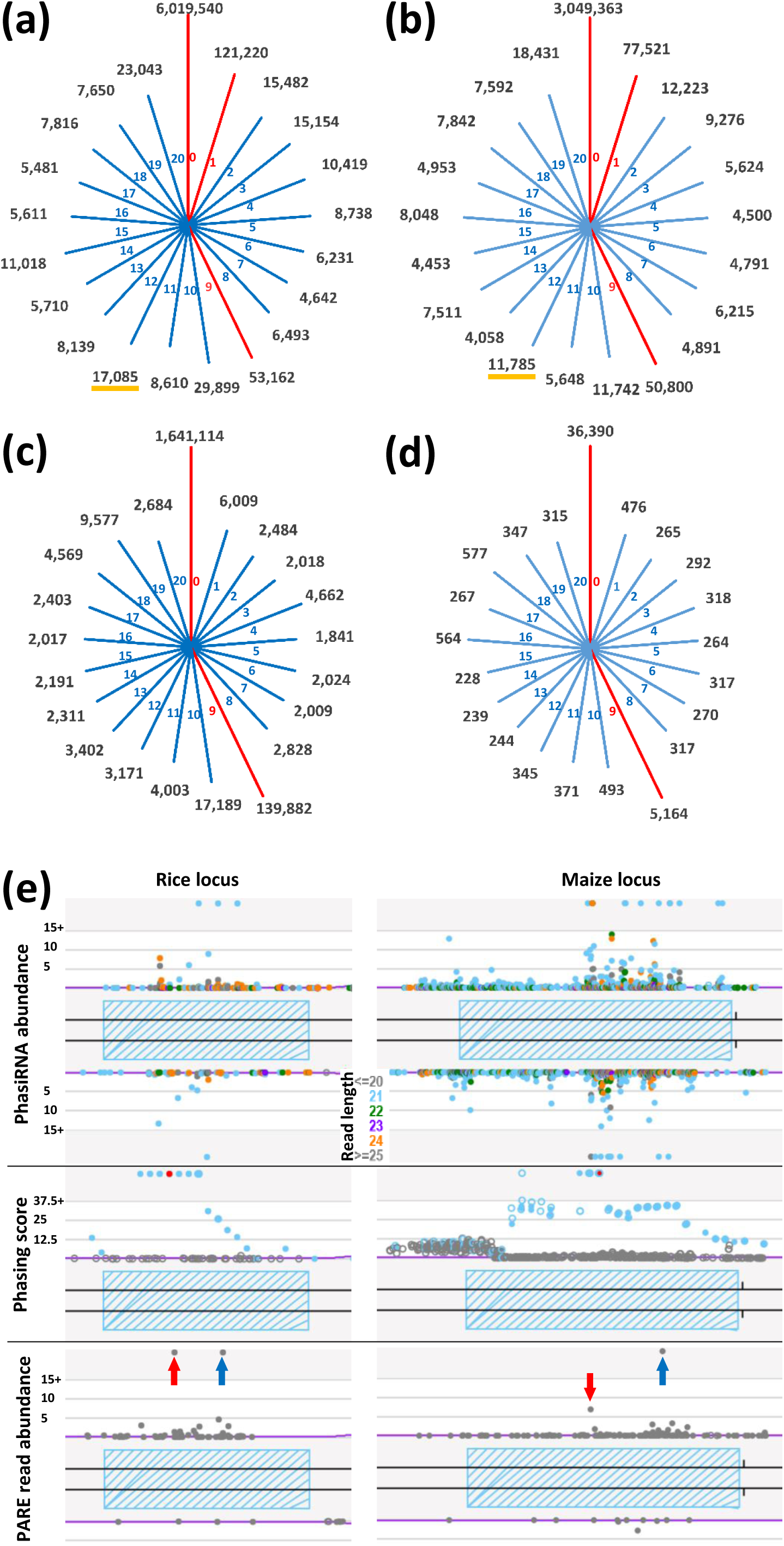
Secondary peaks of 21-nt phasiRNA and PARE data demonstrate *cis* activity. PhasiRNA abundances across 21 registers, starting from the miR2118 cleavage site (register 0) in (a) Stage 4 panicles of rice and (b) 0.4 mm staged anthers of maize. Distribution of Parallel Analysis of RNA ends (PARE) reads across the 21 registers in (c) rice and (d) maize. In each panel from a to d, abundance is represented in transcripts per 10 million (TP10M), hit-normalized (see Methods), and displayed in a log 10 scale. The red lines indicate high abundance registers. The orange underline indicates the 12^th^ register, discussed in more detail in the text. (e) Example of an individual rice (left) or maize (right) phased small interfering RNA (phasiRNA) generating (*PHAS*) locus (cross-hatched box) showing small RNA (sRNA) sizes (colors in key) and abundances (Y-axis, upper panels), phasing scores (Y-axis, middle panels; red dot is the maximum scoring position) and PARE reads (lower panel) as viewed from the Meyers lab website (see Methods). The phasiRNA abundance cutoff is set to 15 for proper display on the website. Red arrows indicate miR2118 cleavage site at register 0; blue arrows indicate cleavage by the phasiRNA corresponding to register 9.

We hypothesized that the peak at register 9 could result from *cis*-cleavage by the premeiotic phasiRNAs. Other 21-nt phasiRNAs such as those from *NB-LRRs* have demonstrated the ability to target and direct cleavage of their own coding transcripts in *cis* (Zhai *et al*. 2011), and thus reproductive 21-nt phasiRNAs might function similarly. We generated PARE data, an approach for genome-scale validation of target cleavage (German *et al*. 2009), from rice anthers and panicles (Supporting Information Table S1) to determine whether the phasiRNAs observed at registers 0 and 9 correspond to cleavage sites. We assigned the PARE reads to registers and observed a strong peak at register 0 corresponding to miR2118 cleavage (Fig. 2c,d). We also observed a peak at register 9, consistent with our hypothesis that cleavage is directed in *cis* by phasiRNAs (Fig. 2c,d). Moreover, visual inspection in genome browsers (see Methods) of individual *PHAS* loci for either rice or maize demonstrated abundant 21-nt phasiRNAs with two peaks in the PARE data (Fig. 2e), one at the site of cleavage directed by miR2118 and a second in the same register (register 9) as cleavage directed by a phasiRNA. This process is discussed in detail below. Thus, these shifts in register are hallmarks of 21-nt phasiRNAs acting in *cis*, directing cleavage of their own precursors.

### Non-templated “tailing” of bottom-strand, first-cycle phasiRNAs

Conventionally understood, the biogenesis of phasiRNAs (and tasiRNAs) produces dsRNA duplexes with a 3’ 2-nt overhang on both termini. This biogenesis includes (i) the synthesis of a complementary ‘bottom’ strand by RDR6 (Rajeswaran and Pooggin 2012), (ii) DCL processing (Elbashir *et al*. 2001), and (iii) stabilization and 3’ protection by HEN1 (HUA ENHANCER1) (Chan *et al*. 2009, Li *et al*. 2005, Yang *et al*. 2006, Yu *et al*. 2005). However, a poorly-described component in this model is the bottom strand of the first phasiRNA; it seems unlikely to be 21 nt, as this would require a 2-nt, single-stranded RNA 3’ end that would require an extension past the miR2118 cleavage site (Fig. S1b). To investigate the size of this bottom-strand, first phasiRNA, we quantified for this position at all 21-*PHAS* loci the sRNAs of 19-, 20-, 21-, and 22-nt and calculated the percentage contribution (other lengths consisted of very few reads and were ignored). The majority of the reads were unexpectedly 20- and 21-nt long (Fig. 3a).

**Fig. 3.**
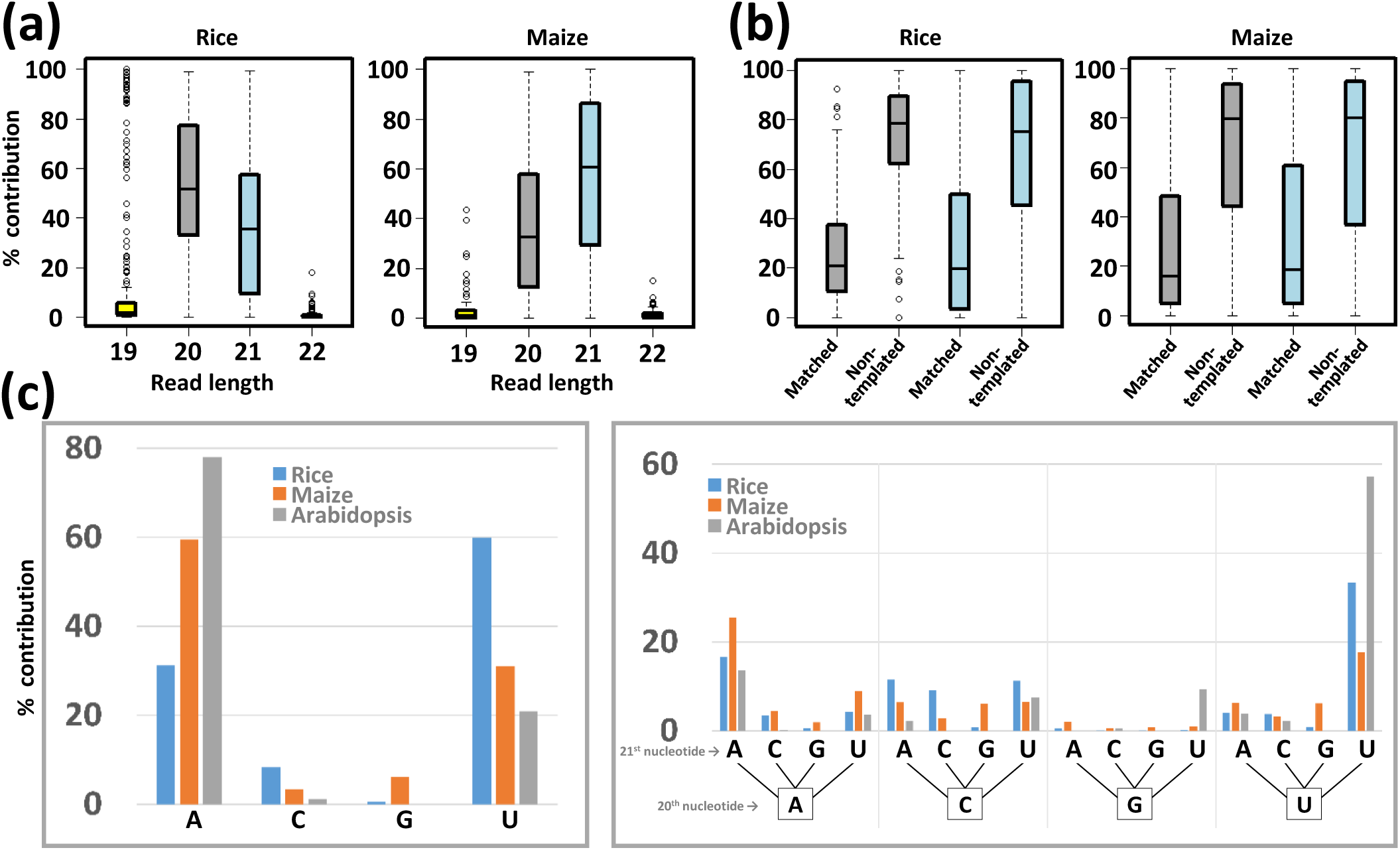
The first bottom-strand phasiRNA undergoes non-templated 3’ modifications. (a) Percentage contribution of 19-, 20-, 21-, and 22-nt reads mapping to the first bottom-strand phased small interfering RNAs (phasiRNAs) in rice and maize. For 20-, 21-, and 22-nt reads, the mapping was constrained only to the first 19 nt. (b) Percentage contribution of “matched” and “non-templated” 20-(colored gray) and 21-nt (colored blue) reads mapping to the first bottom-strand phasiRNAs in rice and maize. Non-templated 20- and 21-nt reads have a 1 or 2-nt 3’ mismatch, respectively; matched reads are complementary at the 3’-most 1 or 2-nt positions, and may reflect either template addition or non-templated tails that happen to match the 3’ fragment (Fig. S1b). (c) Mono-(left) and di-nucleotide (right) percentage contributions of the non-templated 20- and 21-nt reads mapping to the first bottom-strand phasiRNAs in rice and maize, and the same position for *TAS1b*/*c and TAS2* trans acting small interfering RNAs (tasiRNAs) in Arabidopsis.

We next sought to address whether the additional 3’ nucleotides of bottom-strand, first cycle phasiRNAs are “templated” or “non-templated”. To explain this concept, prior work shows that HEN1 SUPPRESSOR 1 (HESO1) and UTP:RNA URIDYLTRANSFERASE 1 (URT1) function in sRNA turnover by uridylating the 3’ termini of unmethylated sRNAs (Ren *et al*. 2012, Tu *et al*. 2015, Zhao *et al*. 2012); this is a non-templated addition (or “tailing”) resulting from terminal transferase activity. Non-templated nucleotides are also observed at the 3’ end of heterochromatic siRNAs (hetsiRNAs) (Wang *et al*. 2016). Biochemical analysis demonstrates that RDR6 has both primary activity as a template-based RNA polymerase as well as activity as a terminal nucleotidyltransferase to add non-templated 3’ nucleotides (Curaba and Chen 2008). We examined the 20- or 21-nt bottom-strand first-cycle phasiRNAs, looking for either perfect complementarity (“templated”) or mismatched complementarity (“non-templated”) in the last two 3’ end positions (positions 20 and 21), relative to the last two positions of the 5’ fragment of the phasiRNA precursor (Fig. S1b). In other words, complementarity at these two nucleotides would match the portion of the miR2118 target site removed after AGO slicing; the other nineteen 5’ positions are complimentary to the first (5’-most) top-strand phasiRNA.

In both rice and maize for this first-cycle, bottom-strand phasiRNA, we noticed a significant percentage of 20- and 21-nt reads possess non-templated nucleotides in the 3’ end (Fig. 3b), suggesting 3’ modification of this phasiRNA via terminal transferase activity. In Arabidopsis, 3’ tailing of sRNAs is strongly biased towards uridylation and adenylation (Li *et al*. 2005, Tu *et al*. 2015, Wang *et al*. 2016). We performed mono- and di-nucleotide analysis to characterize the 3’ additions, adding in published Arabidopsis data (Tu *et al*. 2015) to compare terminal transferase bias between bottom-strand first-cycle tasiRNAs in Arabidopsis with those of 21-nt phasiRNAs in rice and maize. For the mono-nucleotide analysis, we saw a strong bias towards “A” and “U” as an additional non-templated nucleotide across all three species (Fig. 3c). Similarly, for di-nucleotide analysis, we saw a strong bias towards “AA” and “UU” for the additional two non-templated nucleotides (Fig. 3c). Therefore, this bottom-strand, first-cycle phasiRNA is subject to non-templated, terminal transferase activity that extends this phasiRNA to 20 or 21 nucleotides.

### The first-cycle phasiRNA is highly underrepresented in AGO5/MEL1-loaded phasiRNAs

We examined the first-cycle phasiRNAs in detail, assessing the abundance in our libraries relative to the next 19 cycles (20 cycles total x 21 nt = 420 nt of the precursor). This first-cycle position should be most abundant in register 0, as the phasiRNA initiates at the miR2118 cleavage site, and there is no opportunity for shifts in the register. Comparing the total phasiRNA abundance for each cycle across the rice and maize *PHAS* loci, we observed that cycle 1 has a very low abundance relative to subsequent cycles, a “nonstoichiometric” abundance which is unexpected given that the biogenesis should produce consistent abundances across the precursor (Fig. 4a). We hypothesized that the low abundance in cycle 1 could be due to the combination of RDR6-dependent effects on the bottom-strand phasiRNA (i.e. length variation and 3’ modifications, as described above). However, another possible cause of bias against this first-cycle phasiRNA may result from sequence constraints due to the overlap of this duplex with the miR2118 target site (Fig. S1b).

**Fig. 4.**
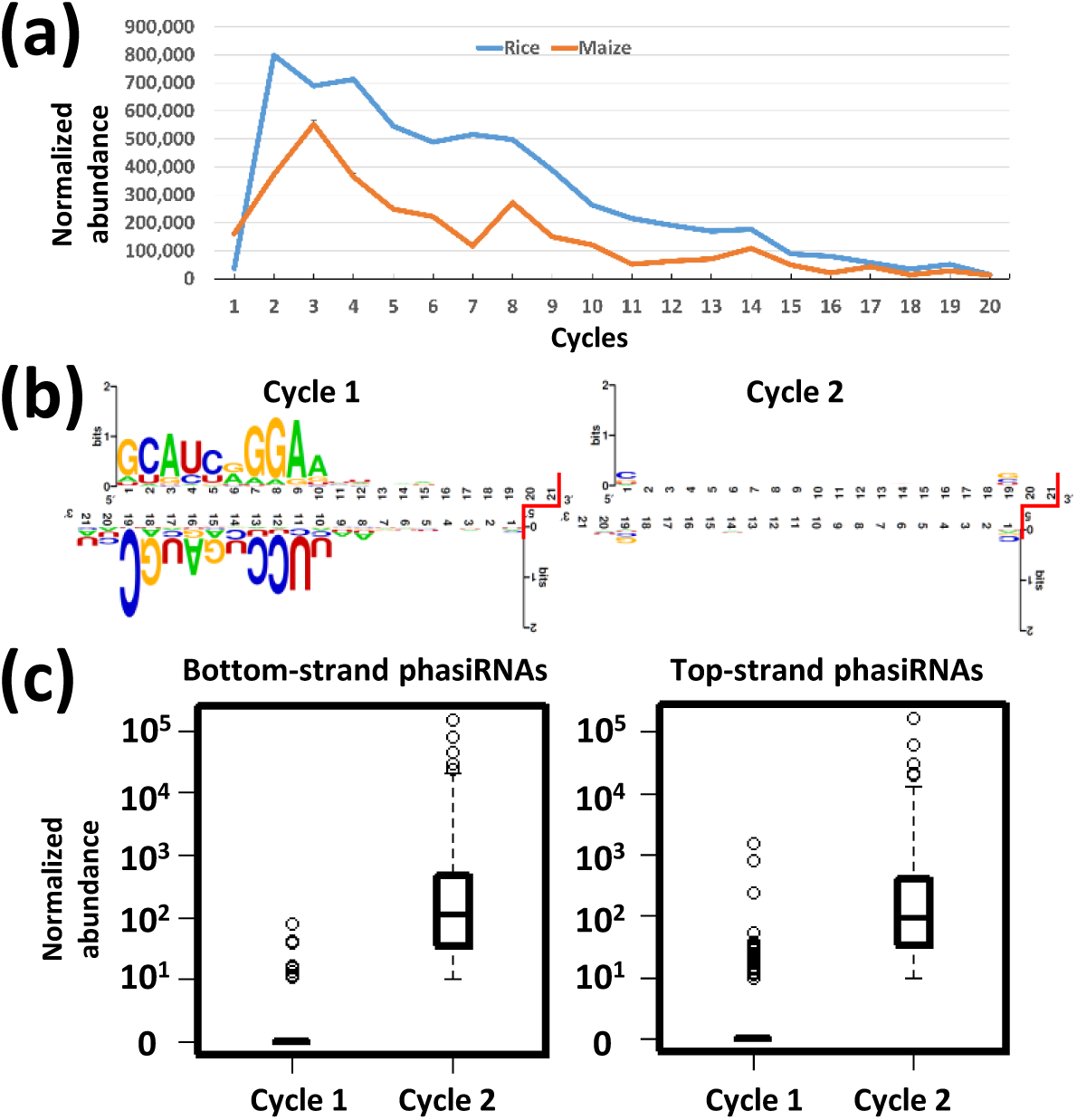
MEL1/AGO5-loaded phasiRNAs show a significant reduction in first-cycle phasiRNAs. (a) Abundance distribution of 21-nt phased small interfering RNAs (phasiRNAs) in rice (blue line) and maize (orange line), measured for twenty cycles downstream (3’) of the miR2118 cleavage site. (b) Sequence motif of top- and bottom-stranded phasiRNAs in cycle 1 (left) and cycle 2 (right) in rice. The connector symbols in red signify the 3’ overhang resulting from Dicer processing. (c) MEIOSIS ARRESTED AT LEPTOTENE 1 (MEL1)-loaded phasiRNAs mapping to cycle 1 (n=11) and cycle 2 (n= 556) of the bottom-strand phasiRNA generating (*PHAS*) locus (left) and to cycle 1 (n=29) and cycle 2 (n=680) of the top-strand *PHAS* locus (right).

To characterize the composition of this phasiRNA duplex, we compared phasiRNAs that mapped to cycle 1 against phasiRNAs that mapped to cycle 2 for both top- and bottom-strands. For bottom-strand reads in cycle 1, we considered all 21-nt reads comprised of 19 perfectly-matched 5’ nucleotides but allowing a 2-nt mismatch at the 3’ end. The consensus sequence showed a strong bias for positions 1 to 10 in the top strand (including a 5’ G), and positions 10 to 19 in the bottom strand, in both rice and maize (Fig. 4b, left panel). We found no bias in the remainder of the phasiRNA duplex. Repeating the analysis for cycle 2 phasiRNAs, the only observed bias was an over-represented cytosine for the first (5’) nucleotide of both strands (Fig. 4b, right panel). Prior work has demonstrated that the first nucleotide of an sRNA influences AGO loading (Mallory and Vaucheret 2010, Mi *et al*. 2008). To assess the impact of this 5’ bias in the first cycle of 21-nt phasiRNAs, we used RNA immunopurification (RIP) data generated from Komiya *et al*. (2014); in rice, 21-nt reproductive phasiRNAs are bound to MEL1 (in the AGO5 family) and are enriched for phasiRNAs with a 5’ C (Komiya *et al*. 2014). We found that the rice MEL1-bound phasiRNAs from cycle 2 are clearly far more abundant than those from cycle 1 (Fig. 4c). Moreover, the phasiRNAs from cycle 2 were from more *PHAS* loci and were of significantly higher abundance compared to those from cycle 1 (Fig. 4c). Therefore, the strong underrepresentation of this first phasiRNA duplex may result from a combination of a constrained 5’ nucleotide (at least on the top strand), the compositional bias due to the overlap with the target site, and the above-described bottom-strand size variation, all of which may negatively impact AGO loading.

### Reproductive phasiRNAs display wide variations in abundance within each *PHAS* locus

As a final analysis, we focused on the abundances of the 21-nt reproductive phasiRNAs within each locus, to assess the relative proportion represented by the top vs bottom strands, or by individual phasiRNAs. The null hypothesis was that all phasiRNA duplexes in a given locus are produced in stoichiometrically equal levels due to the combined activities of RDR6 and DCL4. Any deviation from this equality may result from AGO loading, target interactions, or other sources of differential stabilization of the sRNAs. To test this hypothesis, we ranked phasiRNA duplexes (top + bottom strand abundances, combined) produced from each *PHAS* locus based on their abundance and compared the top ten most-abundant phasiRNA duplexes across all the *PHAS* loci. In both rice and maize, these ten phasiRNA duplexes demonstrated different abundances (Fig. 5a,b), and that there is a significant difference between the most abundant phasiRNA and the other nine. We then used the IP data (Komiya *et al*. 2014) and repeated the same analysis on MEL1-loaded phasiRNAs to see whether phasiRNAs in an AGO protein demonstrate this same difference in accumulation. Indeed, the MEL1-loaded phasiRNA duplexes demonstrated the same pattern observed with rice and maize data (Fig. 5c). From these findings, we concluded that there is a strong accumulation bias within individual *PHAS* loci, such that one single phasiRNA (or the duplex) is much more abundant than those from other cycles.

**Fig. 5.**
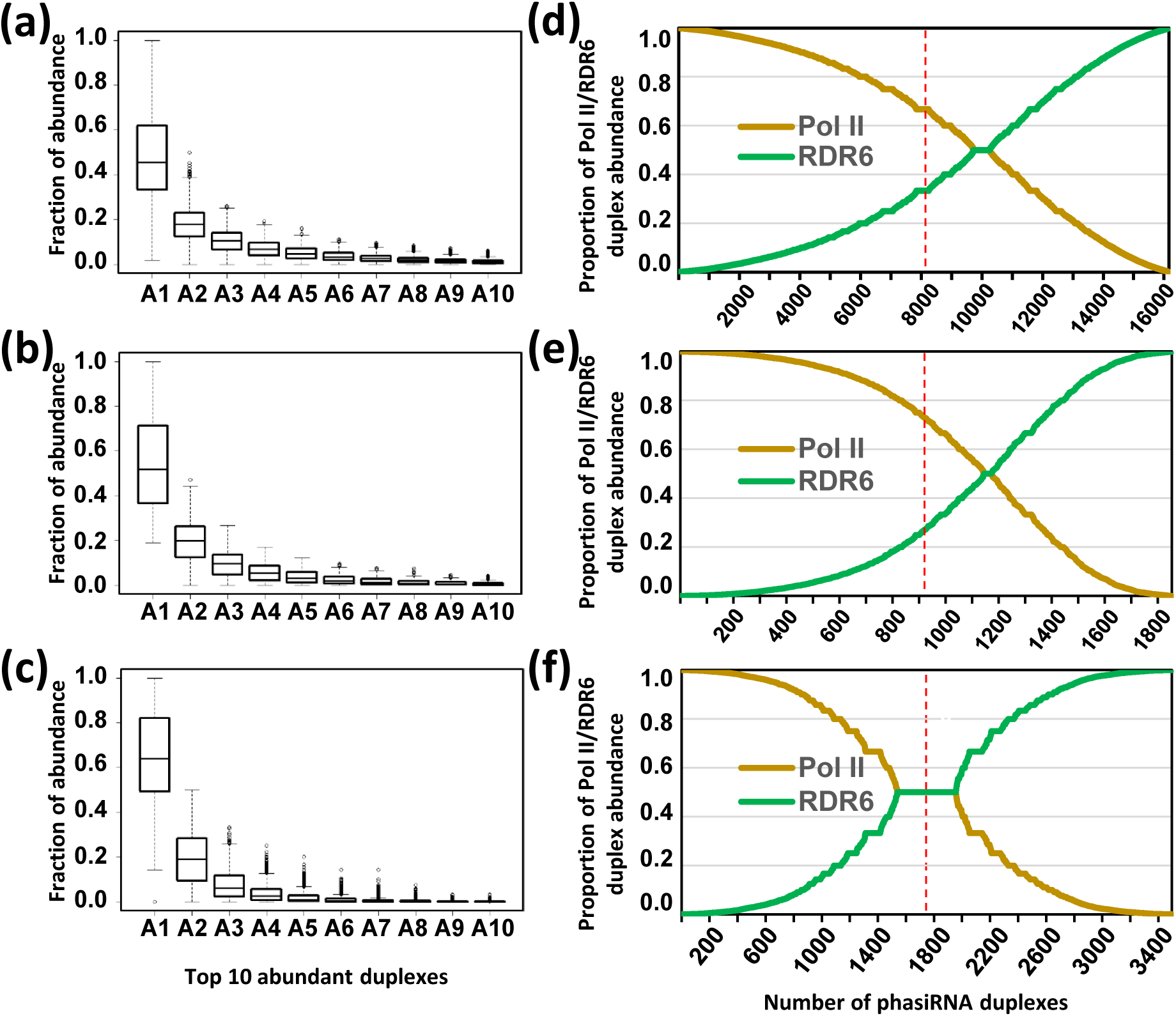
phasiRNAs demonstrate positional and strand biases in abundance within *PHAS* loci. Ranking of the 10 most abundant phased small interfering RNA (phasiRNA) duplexes (A1, most abundant to A10, least abundant) within a phasiRNA generating (*PHAS*) locus and across the *PHAS* loci identified in rice (a) and maize (b), displayed as a fraction of an individual duplex abundance out of the sum of abundance for ten duplexes for the same locus. The duplexes are sorted in descending order of the proportion of the top (RNA POLYMERASE II, Pol II-derived) strand abundance. (c) As in a and b, in this case, the rice MEIOSIS ARRESTED AT LEPTOTENE 1 (MEL1)-loaded phasiRNA duplexes were considered. The duplex abundance is the sum of phasiRNA abundances from both strands making up the duplex. In (d) to (f), the line plots depict the proportions of either the top strand (Pol II) and bottom strand (RNA DEPENDENT RNA POLYMERASE 6, RDR6-derived) phasiRNA abundances for each of the ten most abundant phasiRNA duplexes within a *PHAS* locus and across all the *PHAS* loci identified in rice (d), maize (e), and for MEL1-loaded phasiRNAs (f). The total number of phasiRNA duplexes analyzed for rice, maize and MEL1-loaded phasiRNAs are 16162, 1854, and 3496 respectively. The top (Pol II) strand abundance is represented by a gold line while the bottom (RDR6) strand abundance is represented by a green line. The red dotted line indicates the midpoint in terms of the total number of analyzed phasiRNA duplexes.

To build on this analysis, another way of examining bias within the *PHAS* loci is to assess whether there is a strand bias across the duplexes. Again, the null hypothesis was the top- and bottom-strand phasiRNAs in a given locus are produced in stoichiometrically equal levels. We wanted to assess the presence of a detectable bias in terms of which strand is loaded into AGO. As above, we used the ten most abundant phasiRNA duplexes identified in the previous analysis and compared the proportional abundances of top-to bottom-strand phasiRNAs in each duplex in rice, maize, and MELI-associated phasiRNAs (Fig. 5d,e,f). For both rice and maize, the duplexes with highly disproportionate strand specificity were over-represented for the top (Pol II) versus bottom (RDR6) phasiRNAs: i.e. midpoint crosses the top strand at > 0.6 proportion in rice (Fig. 5d) and > 0.7 proportion in maize (Fig. 5e) compared to the bottom strand with proportion < 0.4 in rice and < 0.3 proportion in maize. The same analysis of MEL1-bound phasiRNAs revealed less bias (Fig. 5f). While the reason for this is unclear, one possibility is that because MEL1 preferentially binds the subset of phasiRNAs with C in 5’ end it displays less of a bias relative to the full set of sequenced, abundant phasiRNAs that could have been loaded into other AGOs in a more highly biased manner. In conclusion, our analysis of representation bias in 21-nt reproductive phasiRNAs demonstrates strong biases in the representation (by abundance) of individual duplexes within a *PHAS* locus, and by the strand that accumulates. This is consistent with a hypothesis in which despite the production of tens of thousands phasiRNAs, only a small proportion are stabilized (presumably due to their functionality), leading to nonstoichiometric abundances of individual phasiRNAs within a *PHAS* locus.

## Discussion

miRNAs and siRNAs, independently and together, play essential roles in RNA silencing, a process that is important during plant development. Previous analyses of maize anthers (Zhai *et al*. 2015) and rice inflorescences and anthers (Fei *et al*. 2016, Johnson *et al*. 2009, Song *et al*. 2012b) reported hundreds of 21- and 24-nt reproductive *PHAS* loci, triggered by miR2118 and miR2275, with clear spatiotemporal patterns in the abundances of the phasiRNAs and their triggers. In this work, we more deeply characterized the sRNAs from these *PHAS* loci to identify characteristics of their biogenesis and accumulation. Focusing on the 21-mers, we propose a model to integrate our new observations with prior data (Fig. 6a); that is, after phasiRNA biogenesis, some phasiRNAs can direct cleavage of their own precursor transcripts, creating an offset of either 9 or 12 nucleotides relative to the trigger (miR2118) cleavage site (Fig. 6b,c). This has several implications, one of which is that 21-nt reproductive phasiRNAs are capable of directing cleavage, not previously demonstrated, although not entirely unexpected for this length of sRNA. A second implication is this phasiRNA-directed cleavage can yield tertiary sRNAs, perhaps similar to the *TAS2 3’D6(*−) tasiRNA in Arabidopsis (Chen *et al*. 2007, Howell *et al*. 2007), but with two important differences: (a) most reproductive phasiRNAs are 21-nt not 22-nt (inconsistent with roles as phasiRNA triggers), and (b) the 12-nt shift indicates that some phasiRNAs act on the RDR6-derived bottom strand, and existing models fail to describe how this could yield additional phasiRNAs.

**Fig. 6.**
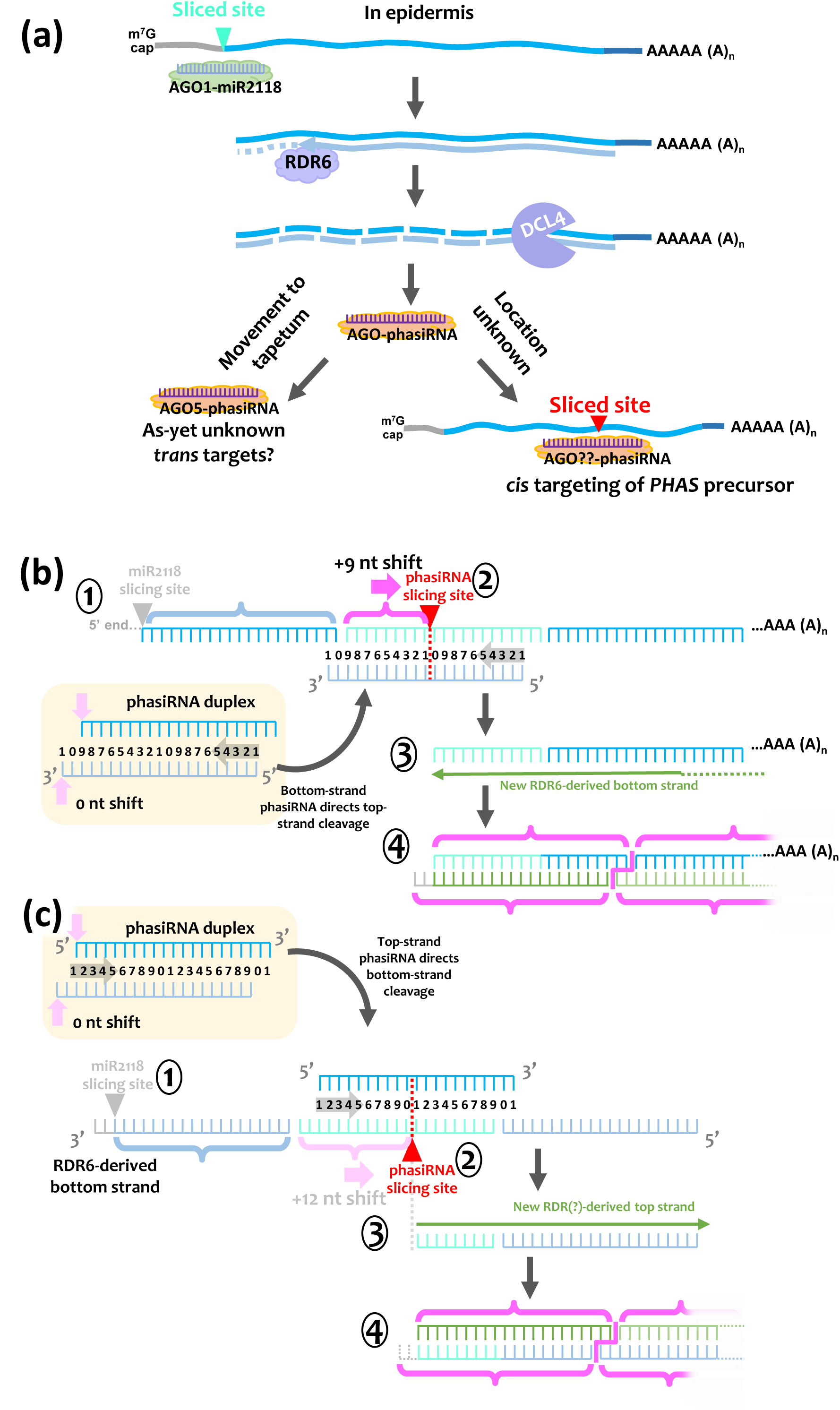
Models of 21-nt phasiRNA activity in *cis*. (a) Overview of 21-nt phased small interfering RNA (phasiRNA) biogenesis. miR2118 directs cleavage of the precursor transcript by ARGONAUTE 1 (AGO1); prior work indicates this occurs in the anther epidermis. RNA DEPENDENT RNA POLYMERASE 6 (RDR6), initiating at an as-yet unclear position in the precursor, synthesizes a complementary strand to convert the cleaved transcript into double-stranded RNA (dsRNA). The dotted line at the terminal end of RDR6 signifies the termination occurs at the sliced top-strand position. DICER-LIKE4 (DCL4) processes the dsRNA, starting from the cleavage site, into 21-nt phasiRNA duplexes (cycles). These 21-nt phasiRNAs are loaded into AGO5, and either direct cleavage of a homologous precursor transcript in *cis* (possibly performed by a different AGO, and in an unknown cell layer), or function in *trans* against as-yet unknown targets, most likely outside of the epidermis. (b) Overview of the process of *cis* targeting by a bottom-strand phasiRNA targeting the RNA POLYMERASE II (Pol II) precursor strand, yielding the observed +9 nt-shifted phasiRNAs. In step 1, the mRNA precursor of phasiRNAs may be either intact or sliced by miR2118 (indicated by grey text). The miR2118 target site designates register 0 initiation of phasiRNAs. In step 2, a bottom-strand phasiRNA targets the messenger RNA (mRNA) precursor, directing slicing between its 10^th^ and 11^th^ nucleotide positions, corresponding to the slicing of the top-strand precursor at the inverted red triangle; this is a +9 nt shift relative to the position typically cleaved within the miR2118 target site. Production of the observed +9 nt-shifted phasiRNAs from this newly sliced mRNA would require recruitment of RDR6 (step 3) and subsequent processing by DCL4 in the 3’ direction (step 4), as is typical for phasiRNA biogenesis. (c) Overview of the process of *cis* targeting by a top-strand phasiRNA targeting the RDR6 precursor strand, yielding the observed +12 nt-shifted phasiRNAs. The top-strand phasiRNA is Pol II-derived and thus has homology only to the RDR6-derived bottom strand (steps 1 & 2). Slicing of this RDR6-derived bottom strand (step 2) must recruit a polymerase to convert the single-stranded RNA to double-stranded (step 3), to serve as a template for DCL4 processing (step 4), yielding the observed tertiary phasiRNAs shifted by +12 nucleotides.

How might these tertiary sRNAs be produced? First, we noticed that the proportion of cleavage products (i.e. PARE reads) in register 9 relative to register 0 was significantly higher (rice: log2 FC = 5.6, p-value < 0.01, maize: log2 FC = 3.6, p-value < 0.01) than the level of sRNAs in register 9 vs register 0 (Fig. 2c,d versus 2a,b); this could indicate that few of the phasiRNA-cleaved precursors are subsequently converted into additional phasiRNAs. In other words, phasiRNA-directed *cis* cleavage might typically occur without subsequent conversion to tertiary sRNAs, consistent with mostly 21-nt phasiRNAs. The smaller proportion of register 9 tertiary sRNAs requires the recruitment of the RDR6-DCL4 machinery (Fig. 6b), perhaps resulting from either (a) the small proportion of phasiRNAs that are 22-nt in length functioning as triggers, or (b) multiple 21-nt phasiRNAs functioning to recruit the machinery in a manner analogous to two-hit tasiRNA biogenesis (Axtell *et al*. 2006). Second, the register 12 phasiRNAs are unexpected: for their biogenesis, AGO-loaded phasiRNAs must be acting on an accessible (ssRNA) RDR6 template (Fig. 6c), yet existing models have the RDR6 and Pol II phasiRNA precursor strands strongly annealed. In addition, to obtain phasiRNAs from a sliced RDR6-derived strand would require a mechanism for biogenesis of a new top strand, yielding a dsRNA that can be processed by DCL4 (Fig. 6c). This top strand, synthesized off of a cleaved, RDR6-derived bottom strand, would require either an additional round of RDR transcription or as yet-undescribed processes or polymerase activities.

The first or cycle 1 phasiRNA duplex is subject to several influences that are unique to this duplex. For example, we observed that the first bottom-strand phasiRNA captured in sequencing libraries is mostly 20- or 21-nt, and not 19-nt as expected. This was explained by a conserved mechanism of small RNA tailing. This mechanism, common between tasiRNAs in Arabidopsis and phasiRNAs in grasses, involves HESO1 and URT1 (Tu *et al*. 2015). We hypothesize that the bottom-strand, cycle 1 phasiRNA is produced as a 19-nt sequence but this length is highly disfavored and mostly eliminated; a proportion of these tailed to 20- or 21-nt, and these are retained, but not at a frequency high enough to impact the overall abundance of this phasiRNA (relative to subsequent cycles). Another possible explanation for this abundance difference comes from the observation of its reduced incorporation into AGO proteins, based on published IP data. This AGO incorporation bias may result from the biased sequence composition of this cycle 1 phasiRNA - it is comprised of about half of the miR2118 target site, and thus may lack signals necessary for AGO loading. For example, there is an enrichment in 5’ C for AGO-loaded 21-nt reproductive phasiRNAs in rice (Komiya *et al*. 2014), and the top-strand, cycle 1 phasiRNA lacks this attribute.

An additional and unexpected bias that we observed in reproductive phasiRNAs is a top/bottom strand bias. Measured across hundreds of loci, the top or Pol II-derived strand accounted for a higher proportion of abundant phasiRNAs in the sequencing data than the bottom, RDR6-derived strand. The reason for this is unclear, but perhaps the two strands vary in so-called epigenetic modifications of RNA, such as methyl-6-adenosine or other post-transcriptional modifications. The two strands are transcribed in different locations (nuclear versus cytoplasmic), and the ‘bottom’ RDR6-dependent strand is presumably processed by DCL4 almost immediately after its biogenesis, so such changes could be introduced during mRNA maturation. In this scenario, the resulting phasiRNAs might look different to the AGO protein, resulting in differential loading and differential stabilization. The lack of bias that we observed in the AGO5/MEL1-loaded data may reflect a different preference for this particular AGO relative to others.

Many of our results describe ways in which the phasiRNA abundances vary from perfect stoichiometry. In other words, theoretically, the long, double-stranded precursor processed by Dicer should give rise to identical abundances of the individual phasiRNAs, which is clearly not the case in the sequencing data. We observed significant biases in the abundances of phasiRNAs that relate to their cycle (position), and to their derivation from the top or bottom strand of the duplex RNA precursor. Assuming that phasiRNA stabilization results from an interaction with a target (Chatterjee *et al*. 2011, Chatterjee and Grosshans 2009) the observation that a single phasiRNA (or duplex) is overrepresented relative to others is consistent with a hypothesis in which individual reproductive phasiRNAs function with specific targets. Within a *PHAS* locus, those with higher abundances may be functional, while those with lower abundances may be degraded. This is in contrast to a hypothesis in which reproductive phasiRNAs function collectively with generic and non-specific targeting. This scenario in which a single phasiRNA from a *PHAS* locus is “functional” could make 21-nt phasiRNAs comparable to miRNAs, or Arabidopsis *TAS* loci in which just one or two tasiRNAs are functionally relevant. Yet, in anthers, there is a synchronized set of hundreds of tissue-specific *PHAS* loci, so clearly the activity of many phasiRNAs is required at a particularly moment in development. We note that SNPs at single phasiRNAs in two rice 21-nt *PHAS* loci are the genetic basis for the agronomically-important photoperiod-sensitive male sterility phenotype (Ding *et al*. 2012, Fan *et al*. 2016, Mei *et al*. 1999, Zhu and Deng 2012). With these points in mind, future studies should examine the conservation of abundances and targets of individual phasiRNAs across orthologous *PHAS* loci in diverse grass species.

## Acknowledgements

Work on this project in the Meyers lab is supported by NSF Plant Genome Research Program award # 1339229. Work in the Zhang lab is supported by the National Natural Science Foundation of China (NSFC award # 91540101). Work in the Zhai lab is supported by the Thousand Talents Program for Young Scholars and by the Program for Guangdong Introducing Innovative and Entrepreneurial Team (2016ZT06S172). We thank members of the Meyers and Zhang labs for helpful discussion.

## Authors’ contributions

S.T., Z.C., and B.C.M. designed the research. Z.C. collected the samples, Z.C., S.M., and J.Z. prepared the libraries. S.T. conducted the analysis. S.T., Z.C., and B.C.M. wrote the manuscript. All authors reviewed and approved the final manuscript.

**Supporting Information: Table S1**. Summary of small RNA sequenced libraries.

**Supporting Information: Table S2**. List of identified 21- and 24-nt phased small interfering RNA (phasiRNA) generating (*PHAS*) loci.

**Supporting Information: Fig. S1**. Illustration of cycles and registers in phased small interfering RNA (phasiRNA) biogenesis.

In this work, and here focusing on just 21-nt phasiRNAs, we extensively use the terms “register” and “cycle” to describe the positions of phasiRNAs relative to the target site of the miRNA trigger. A register is measured in single nucleotide increments relative to the position of the cleavage site initiated by the miRNA trigger, analogous to a reading frame in an mRNA; therefore, for reproductive phasiRNAs, the register resets every 21 nucleotides, and if a phasiRNA precursor is perfectly processed (phasiRNAs of exactly the same length), all the phasiRNAs will be in the same register. A cycle is a unit of a phasiRNA, analogous to a codon within an mRNA, measured in units each of length 21 nt; there are an indeterminate number of cycles per precursor, limited only by the length of the transcript 3’ to the miRNA trigger site. These concepts are illustrated in this figure. (a) DICER-LIKE4 (DCL4) processes the cleaved precursor in cycles of 21-nt, starting from the miR2118 cleavage site. For our analyses, twenty cycles starting from the miR2118 cleavage site were considered. (b) Zoomed-in view of processed transcript: the inverted green triangle corresponds to the miR2118 cleavage site and this nucleotide position is designated as register 0; all the cycles that follow, spaced by exactly 21 nucleotides, are also in register 0. The small red box indicates the two nucleotides in the first, bottom-strand phasiRNA that our analyses indicate are added by terminal transferase modifications (“tailing”). (c) Depiction of register 1; this corresponds to a 1-nt shift to the 3’ direction (to the right) relative to the miR2118 cleavage site, as compared to no shift in b. Such shifting therefore defines 21 “registers” (0 to 20 nt) in phasiRNA positions relative to the initial miR2118 sliced site.

