## Supplementary figures and images for "*Cis*-directed cleavage and nonstoichiometric abundances of 21-nt reproductive phasiRNAs in grasses"

### Supplementary Materials

Tamim et al., Figure S1

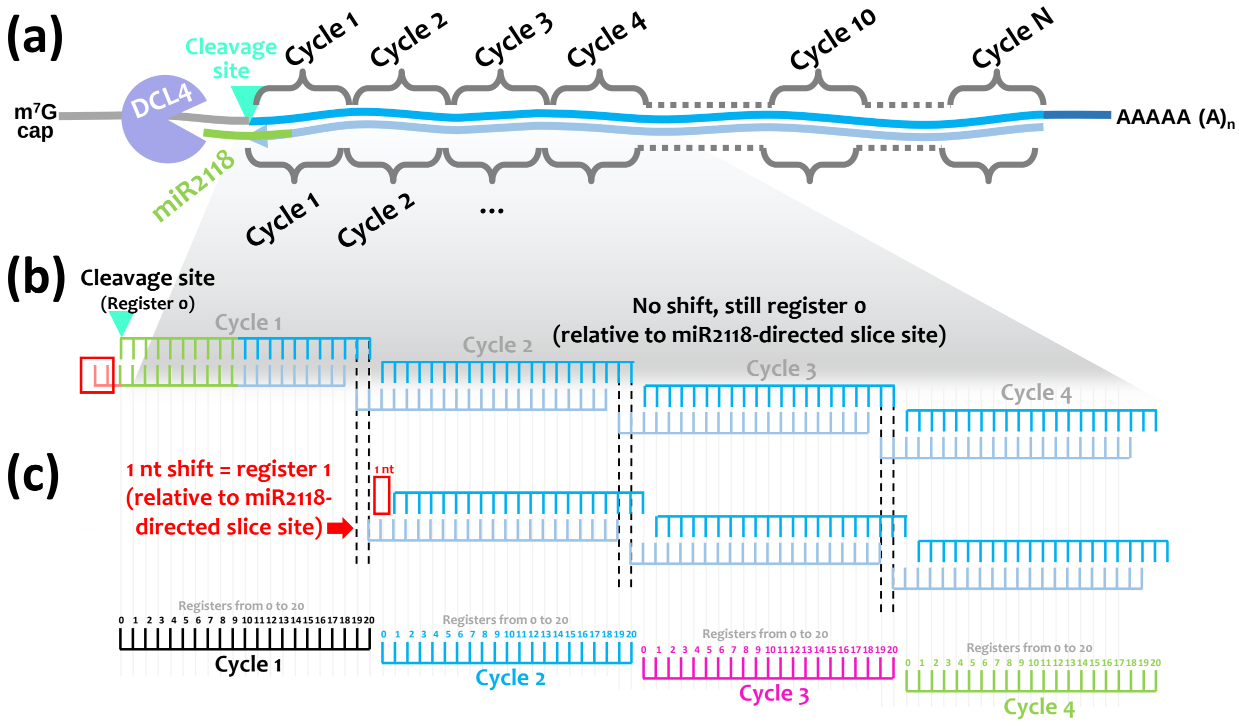
